# Relationship between prokaryotic GC content and environmental salinity

**DOI:** 10.1101/2023.05.07.539728

**Authors:** En-Ze Hu, Shen Sun, Deng-Ke Niu

## Abstract

**Background:** The correlation between GC content and halophilicity has received limited attention, despite the numerous environmental factors associated with GC content evolution. While higher GC content has been linked to halophiles in some archaeal cases, it is widely believed that selective pressure from high-intensity ultraviolet radiation in halophilic archaea drives GC content increase, as it prevents DNA photoproduct formation. However, this assumption has not been statistically analyzed in a phylogenetically independent manner prior to our study.

**Results:** Using phylogenetic generalized least squares, we investigated the relationship between GC content and halophilicity in 1226 bacteria and 181 archaea. Our analysis found significant positive correlations in bacteria but not in archaea. Resampling analysis indicates that the absence of significant correlation in archaea may be due to the relatively small sample size. We also observed that the strength of the correlation is negatively influenced by the functional constraint of genomic components. Additionally, we found that halophilic bacteria and archaea do not have lower photoreactivity (a measure of DNA vulnerability to ultraviolet radiation) than the photoreactivity expected from their GC contents.

**Conclusions:** In contrast to previous assumptions, we did not find evidence to support the widespread photoprotection hypothesis or another hypothesis that high GC content in halophiles stabilizes nucleic acid structures. Instead, our findings align with a nonadaptive hypothesis. Halophilic prokaryotes likely evolved high GC content due to frequent GC-biased gene conversion in response to DNA double-strand breaks induced directly or indirectly by high salt concentrations.

## Background

GC content refers to the proportion of guanine and cytosine present in nucleotide sequences. This fundamental characteristic stabilizes the double-stranded DNA helix and three-dimensional RNA structures. It can affect various aspects of genetic information performance, including three-dimensional genome organization (1), mutation and recombination rates (2), gene transcription and mRNA translation rates (3-5), spicing site recognition (6), RNA interference efficiency (7), and horizontal gene transfer (8, 9). Many environmental factors, such as growth temperature, nutrient availability, and ultraviolet (UV) radiation (10-17), have been found or proposed to be associated with the evolution of GC content.

A wide range of high-salinity environments can be found globally, from naturally occurring hypersaline lakes and deep-sea brines to artificially created salterns. Halophiles have been discovered in all three domains of life, including Archaea, Bacteria, and eukaryotes (18). The first sequenced halophilic archaeon, *Halobacterium* sp. NRC-1, comprises a primary chromosome and two microchromosomes with GC content of 68%, 58%, and 59%, respectively (19). Early studies suggested that high GC content is a characteristic feature of extreme halophiles (20-22). However, *Haloquadratum walsbyi*, an extremely halophilic organism, has a low GC content of 47.9% (23). In a later study, Jones and Baxter (24) compared the GC content of 29 halophilic archaea and 2231 other prokaryotes using a Welch two-sample t-test and observed a remarkably significant difference (63.1% ± 1.3% vs. 49.7% ± 0.55%, *P* < 2.2 × 10^−16^). However, they did not consider the phylogenetic non-independence among species. Comparative analysis of phylogenetically non-independent data using traditional statistical methods often produces erroneous results (25-27). The phylogenetic non-independence of halophilic archaea is of particular concern because they are concentrated in a few phylogenetic lineages (18, 28).

To confidently determine the relationship between GC content and halophilicity, we conducted phylogenetic generalized least squares (PGLS) regression analyses (29) on 1226 bacterial species (816 non-halophilic and 410 halophilic) and 181 archaeal species (8 non-halophilic and 173 halophilic). Additionally, we explored the potential mechanisms underlying the correlation between GC content and halophilicity and revisited the prevalent hypothesis of avoiding UV-induced thymidine dimer formation (30) using the current dataset and phylogenetic comparative methods.

## Results

Prokaryotes are categorized into four groups based on their optimal range of salt concentration for growth: non-halophiles, slight halophiles, moderate halophiles, and extreme halophiles (31). To obtain a dataset for halophilicity, we utilized the Halodom database (32), Protrait database (33), and Madin et al. prokaryotic trait dataset (34). For PGLS regression analysis, halophilicity was transformed into an ordered categorical variable that assigned numerical values to the four categories (1, 2, 3, and 4).

We re-annotated the representative genomes of the prokaryotic species (identified as reference or representative genomes in the NCBI Genome Database). Considering the strength of functional constraints, we divided prokaryotic genomes into seven genomic components and calculated their GC content: the first and second codon positions of protein-coding sequences (GC_1+2_), the third codon position of protein-coding sequences (GC_3_), intergenic sequences (GC_inter_), tRNA genes (GC_tRNA_), 5S rRNA genes (GC_5S_), 16S rRNA genes (GC_16S_), and 23S rRNA genes (GC_23S_) (Additional file 1: Table S1-S2). It should be noted that the sample size for rRNA genes might be slightly smaller due to incomplete annotations in some genomes.

### Both genomic GC content and halophilicity have strong phylogenetic signals

Horizontal gene transfer among prokaryotes makes genealogical relationships somewhat complex. If bifurcation is not the dominant pattern of phylogeny and common ancestors do not significantly influence the trait values of their descendants, conventional statistical methods may be used. To test whether phylogenetic comparative methods are necessary, we calculated Pagel’s λ values to measure the correlation between the evolution of the analyzed trait and the presumed phylogenetic tree (35). Our analysis revealed that the GC contents of whole genomes and genomic components of both bacteria and archaea in our dataset show strong phylogenetic signals (Additional file 1: Table S3). Similarly, the halophilicity of bacteria and archaea displays significant phylogenetic signals (Additional file 1: Table S3).

Halophilic archaea are naturally exposed to high-intensity UV radiation, and their high GC content was initially thought to be an adaptation that prevents the formation of UV-induced thymine dimers (19, 20, 30, 36). However, further analysis revealed that all four adjacent pyrimidines (TT, TC, CT, CC) are susceptible to photodamage under UV irradiation (37). To quantify photoreactivity (PR), Jones and Baxter (24) developed a formula that takes into account the frequencies of adjacent pyrimidines susceptible to photodamage under UV irradiation (TT, TC, CT, CC) and their intrinsic photoreactivity: PR = 1.73(qTC) + 1.19(qTT) + 0.61(qCT) + 0.39(qCC), where qTC, qTT, qCT, and qCC represent the frequencies of each adjacent pyrimidine. GC content influence DNA photoreactivity since it affects the frequencies of the four adjacent pyrimidines. Our analysis of the photoreactivity of whole genomes and genomic components of bacteria and archaea in our dataset (Additional file 1: Table S4-S5) also revealed strong phylogenetic signals (Additional file 1: Table S3).

### Whole genome GC content significantly correlates with halophilicity in bacteria but not archaea

To investigate this relationship, we used halophilicity as the independent variable and GC_w_ as the dependent variable in PGLS regression analyses. Regression slopes with a plus sign indicate a positive correlation, while those with a minus sign indicate a negative correlation. We used four commonly used models: Brownian motion, Ornstein-Uhlenbeck, Pagel’s λ, and Early burst. The best-fit model was determined by selecting the model with the smallest Akaike information criterion (AIC) value. As shown in Table 1, the Pagel’s λ model is the best fit for both bacteria and archaea, with considerably smaller AIC values compared to the other models. Under this model, we found a significant positive correlation between the genome-wide GC content and halophilicity in bacteria (*P* = 0.015) but not archaea (*P* = 0.908).

**Table 1.**
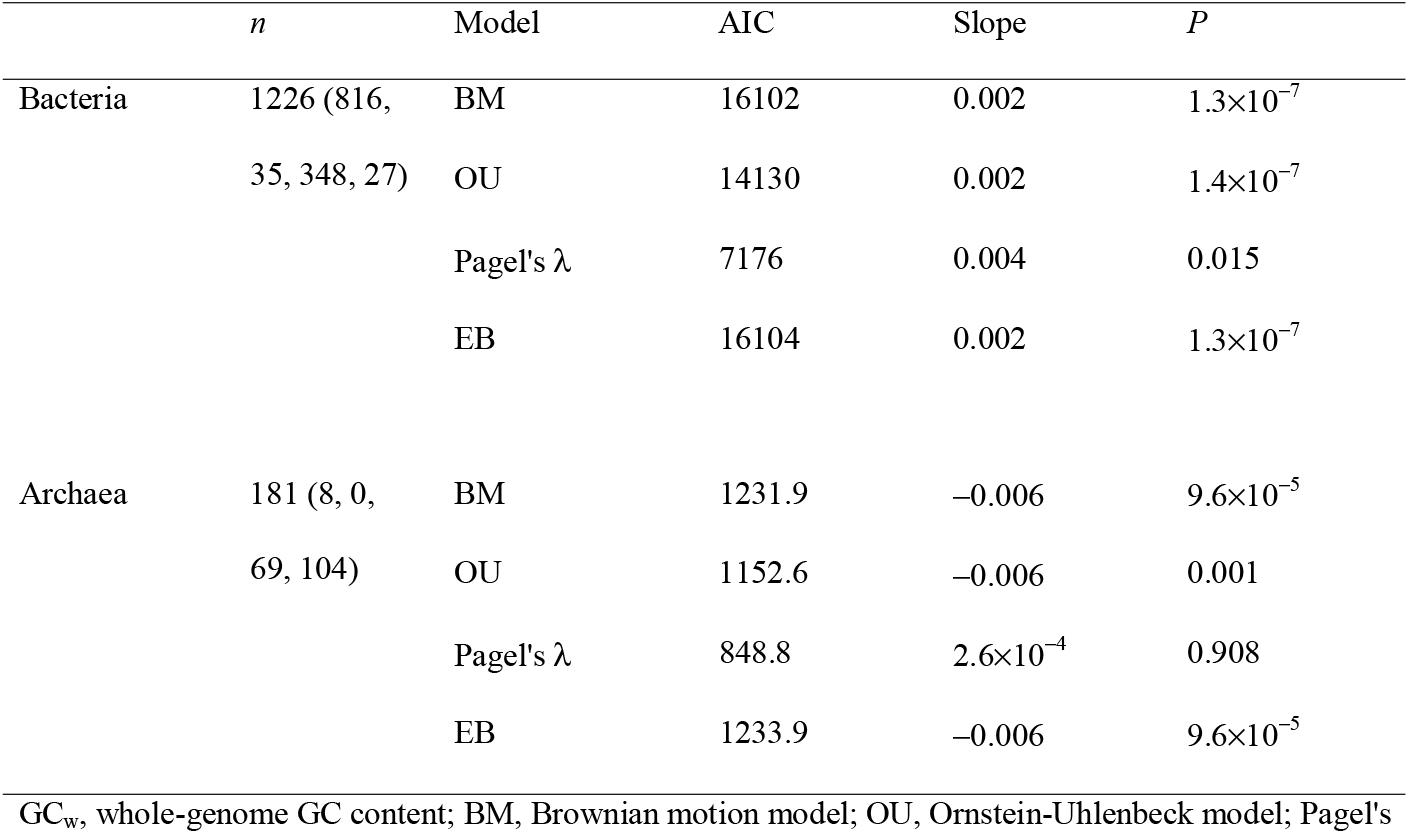
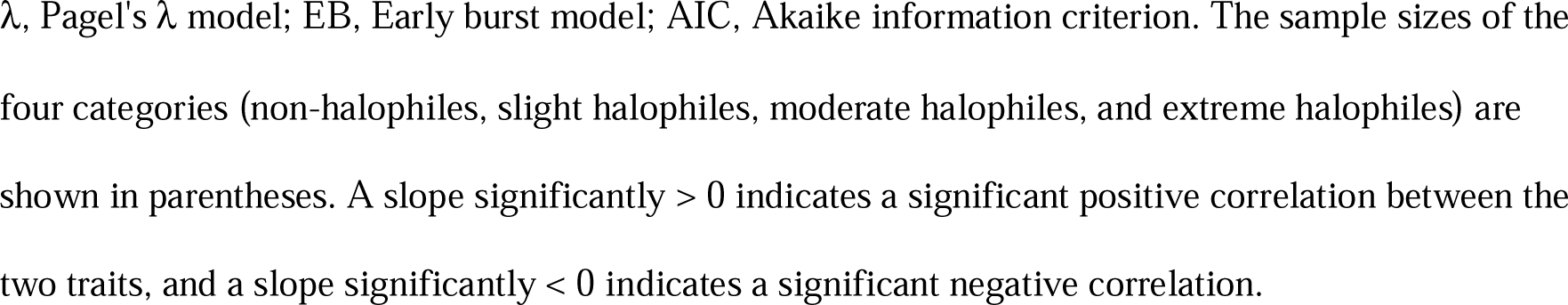
PGLS regression analyses of whole-genome GC content and halophilicity

|  | *n* | Model | AIC | Slope | *P* |
| --- | --- | --- | --- | --- | --- |
| Bacteria | 1226 (816, 35, 348, 27) | BM | 16102 | 0.002 | $1.3 \times 10^{-7}$ |
| | | OU | 14130 | 0.002 | $1.4 \times 10^{-7}$ |
| | | Pagel's $\lambda$ | 7176 | 0.004 | 0.015 |
| | | EB | 16104 | 0.002 | $1.3 \times 10^{-7}$ |
| Archaea | 181 (8, 0, 69, 104) | BM | 1231.9 | -0.006 | $9.6 \times 10^{-5}$ |
|  |  | OU | 1152.6 | -0.006 | 0.001 |
| | | Pagel's $\lambda$ | 848.8 | $2.6 \times 10^{-4}$ | 0.908 |
| | | EB | 1233.9 | -0.006 | $9.6 \times 10^{-5}$ |

$GC_w$ , whole-genome GC content; BM, Brownian motion model; OU, Ornstein-Uhlenbeck model; Pagel's
$\lambda$ , Pagel's $\lambda$ model; EB, Early burst model; AIC, Akaike information criterion. The sample sizes of the four categories (non-halophiles, slight halophiles, moderate halophiles, and extreme halophiles) are shown in parentheses. A slope significantly $> 0$ indicates a significant positive correlation between the two traits, and a slope significantly $< 0$ indicates a significant negative correlation.

Closely related lineage exhibit highly similar GC content and halophilicity due to minimal evolutionary changes since their divergence from a common ancestor. Therefore, such lineages should not be considered as independent samples in phylogenetic comparative studies, and the effective sample size should be well below the census number of the analyzed lineages. We suspect that the lack of a significant correlation between GC_w_ and halophilicity in archaea might be attributed to the small sample size. To test this hypothesis, we randomly resampled 181 species from the 1226 bacteria 1000 times. Results of PGLS regression analysis indicated that significant positive correlations between GC_w_ and halophilicity disappeared in most cases (63.7%, *P* > 0.05). If we analyze a bacterial sample as small as the archaeal sample, we might not be able to find a significant correlation between GC_w_ and halophilicity.

### Relationship between halophilicity and GC content of genome components

We also performed PGLS regression analysis on genome components using the four models above, and Pagel’s λ model was consistently preferred due to the smallest AIC value. In bacteria, halophilicity is significantly correlated with GC_3_, GC_inter_, GC_tRNA_, and GC_5S_ (*P* = 0.036, 0.006, 0.001, and 1.50×10^−4^, respectively), and there is a marginal correlation with GC_1+2_ and GC_23S_ (*P* = 0.059 and 0.096, respectively). However, there is no correlation between bacterial halophilicity and GC_16S_ (*P* = 0.432). The significance of the correlation between GC content and halophilicity increases from the first and second codon positions, the third codon position to intergenic sequences (Table 2). Likewise, the GC content of tRNA and 5S rRNA genes is more strongly correlated with halophilicity than the GC content of 16S rRNA and 23S rRNA genes (Table 2).

**Table 2.** Correlations between GC content of genome components and halophilicity in bacteria

|  | *n* | Slope | *P* |
| --- | --- | --- | --- |
| GC<sub>1+2</sub> ~ Halophilicity | 1226 | 0.001 | 0.059 |
| GC<sub>3</sub> ~ Halophilicity | 1226 | 0.007 | 0.036 |
| GC<sub>inter</sub> ~ Halophilicity | 1226 | 0.005 | 0.006 |
| GC<sub>tRNA</sub> ~ Halophilicity | 1226 | 0.001 | 0.001 |
| GC<sub>5S</sub> ~ Halophilicity | 1109 | 0.004 | 1.50×10<sup>-4</sup> |
| GC<sub>16S</sub> ~ Halophilicity | 1083 | 3.1×10<sup>-4</sup> | 0.432 |
| GC<sub>23S</sub> ~ Halophilicity | 1038 | 8.6×10<sup>-4</sup> | 0.096 |

The correlations were examined by PGLS regressions using Pagel's $\lambda$ model. A slope significantly $> 0$ indicates a significant positive correlation between the two traits, and a slope significantly $< 0$ indicates a significant negative correlation. GC<sub>1+2</sub>, GC content of the first and second codon positions of protein-coding sequences; GC<sub>3</sub>, GC content of the third codon positions of protein-coding sequences; GC<sub>inter</sub>, GC content of intergenic sequences; GC<sub>tRNA</sub>, GC content of tRNA genes; GC<sub>5S</sub>, GC content of 5S rRNA genes; GC<sub>16S</sub>, GC content of 16S rRNA genes; GC<sub>23S</sub>, GC content of 23S rRNA genes.

In archaea, we found no significant correlation between halophilicity and GC content of any genome components (*P* > 0.10 in all cases. Additional file 1: Table S6).

### Photoreactivity is not positively correlated with halophilicity

To investigate whether high GC content prevents UV-induced pyrimidine dimers in bacterial genomes (19, 20, 30, 36), we examined the correlation between halophilicity and photoreactivity of different components of bacterial genomes. Our PGLS regression analysis, using halophilicity as an independent variable and photoreactivity as the dependent variable, found no significant correlation between photoreactivity of bacterial whole-genome sequences, protein-coding genes, tRNA genes, 16S rRNA genes, and 23S rRNA genes and halophilicity (*P* > 0.100 for all five cases, as shown in Table 3). However, intergenic sequences and 5S rRNA genes show significant negative correlations with halophilicity (Table 3). We also analyzed archaea and found no significant correlations between halophilicity and photoreactivity (*P* > 0.100 for all cases. Additional file 1: Table S7).

**Table 3.** Relationship between photoreactivity and halophilicity in bacteria

|  | *n* | Slope | *P* |
| --- | --- | --- | --- |
| PR<sub>w</sub> ~ Halophilicity | 1226 | -4.9×10<sup>-4</sup> | 0.220 |
| PR<sub>p</sub> ~ Halophilicity | 1226 | -3.9×10<sup>-4</sup> | 0.336 |
| PR<sub>inter</sub> ~ Halophilicity | 1226 | -0.001 | 0.002 |
| PR<sub>tRNA</sub> ~ Halophilicity | 1226 | -2.4×10<sup>-4</sup> | 0.122 |
| PR<sub>5S</sub> ~ Halophilicity | 1109 | -0.001 | 0.026 |
| PR<sub>16S</sub> ~ Halophilicity | 1083 | -1.8×10<sup>-4</sup> | 0.210 |
| PR<sub>23S</sub> ~ Halophilicity | 1038 | -1.4×10<sup>-4</sup> | 0.349 |

The relationships were examined by PGLS regressions using Pagel's $\lambda$ model. A slope significantly > 0 indicates a significant positive correlation between the two traits, and a slope significantly < 0 indicates a significant negative correlation. PR<sub>w</sub>, the photoreactivity of whole-genome sequences; PR<sub>p</sub>, the photoreactivity of protein-coding genes; PR<sub>inter</sub>, the photoreactivity of intergenic sequences; PR<sub>tRNA</sub>, the photoreactivity of tRNA genes; PR<sub>5S</sub>, the photoreactivity of 5S rRNA genes; PR<sub>16S</sub>, the photoreactivity of 16S rRNA genes; PR<sub>23S</sub>, the photoreactivity of 23S rRNA genes.

In a genome rich in GC, the occurrence of fewer TTs and more CCs could be a random chance. As the formula that quantifies photoreactivity weighs qTT more than qCC (1.19 vs. 0.39), a low photoreactivity in a GC-rich genome could also be attributed to chance. If a low PR is a result of evolutionary adaptation to avoid UV-induced pyrimidine dimers, it should be lower than the expected chance of occurrence. The expected PR of a DNA sequence, E(PR), could be calculated after randomly arranging bases in the sequence (Additional file 1: Table S4-S5). The observed PR is compared with the E(PR) in three datasets (410 bacteria including slight halophiles, moderate halophiles, and extreme halophiles, 375 bacteria including moderate halophiles and extreme halophiles, and 27 bacteria including only extreme halophiles) using phylogenetic paired t-test (38). The results do not show any significant difference in the first two datasets (Table 4). However, in the 27 extremely halophilic bacteria, the PR and E(PR) showed significant differences in the whole-genome sequences, protein-coding genes, intergenic sequences, tRNA genes, 16S rRNA genes, and 23S rRNA genes (Table 4). Upon further examination of the values, it was observed that the PR values of these sequences are higher than the E(PR) values. The same analysis for archaeal genomes also showed no significant differences (*P* > 0.100 for all cases. Additional file 1: Table S8).

**Table 4.**
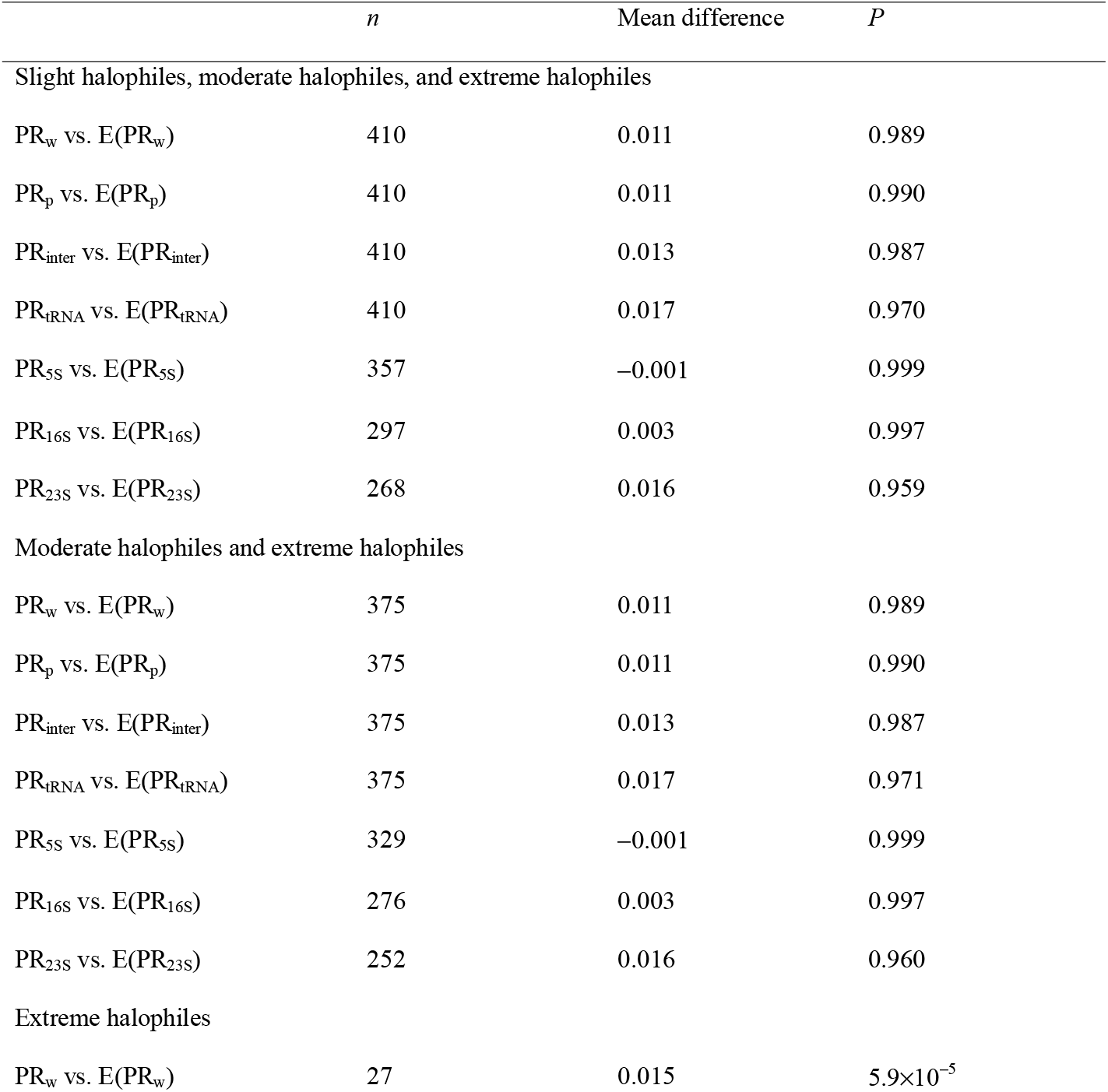
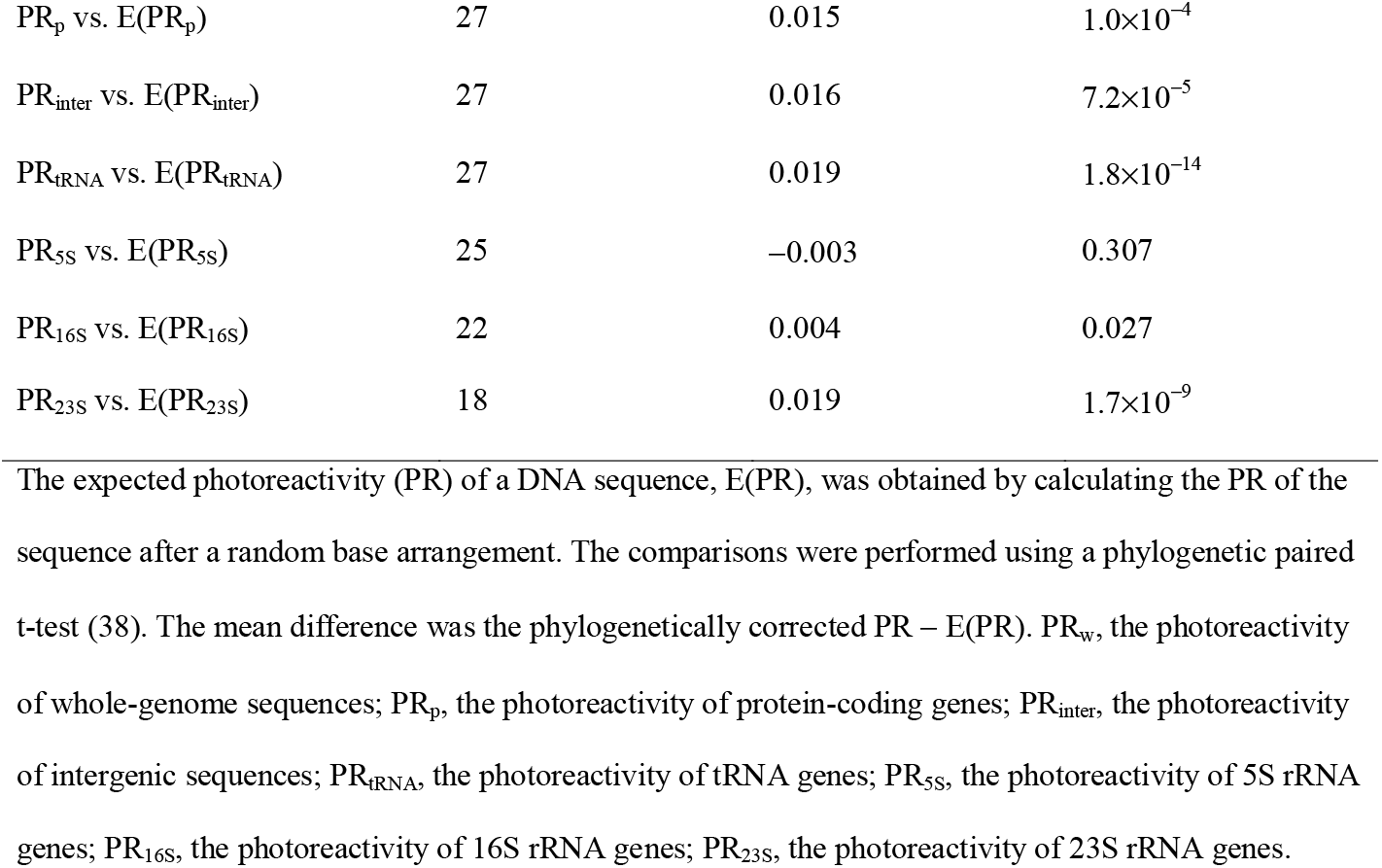
Comparison between observed photoreactivity and the photoreactivity expected by GC content in bacteria

In summary, the results do not support the hypothesis that GC-rich genomes in halophiles evolved to avoid UV-induced pyrimidine dimers.

## Discussion

The relationship between environmental salinity and GC content evolution has not been extensively explored compared to other environmental factors. In the early days of genome sequencing, some halophilic archaea were observed to have remarkably GC-rich genomes (19-22). However, the only previous statistical analysis on the correlation between GC content and halophilicity had a methodological flaw as the phylogenetic non-independence of the prokaryotic data was not considered (24). Our PGLS regression analyses confirmed the correlation in 1226 bacteria but not 181 archaea (Table 1). To address the small sample size of archaea, we randomly resampled 181 species from the 1226 bacteria 1000 times and found that in most cases, the correlations between GC content and halophilicity become statistically nonsignificant. This lack of significant correlation in archaea, similar to our previous study on the relationship between GC content and growth temperature (11), is probably due to the small sample size.

The high concentration of NaCl is typically caused by solar-driven water evaporation in the unique habitats of halophilic archaea, which are typically characterized by high-intensity UV radiation. UV penetration is more substantial in saline water (39), and high salt content in water bodies and coastal areas can affect local atmospheric conditions, reducing ozone concentrations and leading to increased UV radiation (40). Under UV radiation, all four types of adjacent pyrimidines (TT, TC, CT, CC) can potentially undergo photodamage (37). Although high GC content was hypothesized to be an adaptation of the archaea to intense UV radiation (19, 20, 30, 36), the photoreactivity formula (24)[PR = 1.73(qTC) + 1.19(qTT) + 0.61(qCT) + 0.39(qCC)] shows that GC content does not have the strong effect previously assumed in avoiding UV-induced DNA damage. However, photodamage is still a potential driver of GC content evolution, but our analysis in halophilic bacteria or archaea did not find a lower photoreactivity than expected by chance. Therefore, high GC content in halophilic archaea or bacteria should not be interpreted as a photoprotection strategy.

Another possible adaptive effect of high GC content in halophiles is increasing DNA stability under high salt concentrations (41). The low concentrations of cations in the solution can neutralize the negative charge of the phosphate groups on DNA molecules, reducing the repulsive forces between the strands and stabilizing the DNA double helix structure (42, 43). However, when the concentration exceeds a certain threshold, 1M, it can destabilize the DNA double helix structure (44). Some halophilic prokaryotes using the “high-salt-in” strategy need to accumulate molar concentrations of inorganic salts in the cytoplasm, and such salt concentrations exceed the threshold (18, 45). Many halophilic prokaryotes employ the “compatible solute” strategy to cope with osmotic stress by accumulating high concentrations of small organic solutes, such as proline and betaine. These small organic solutes can reduce the stability of the DNA double helix (46-48). The additional hydrogen bonds of G:C base pairs compared to A:T base pairs can help maintain genome stability in high salt concentrations. Therefore, high GC content is also favored in halophilic prokaryotes, whether they accumulate inorganic salts or small organic solutes. However, our genome component observations do not support the DNA double helix stability hypothesis. As DNA double helix structures are more stable than the three-dimensional structures of structural RNA molecules, they are more resistant to destabilizing factors. Therefore, whole-genome GC content correlates less strongly with growth temperature than structural RNA genes (11). However, in this study, we observed that the GC content of 16S and 23S rRNA genes is not significantly correlated with halophilicity (Table 2). Furthermore, the principle of natural selection does not support the DNA double helix stability hypothesis. A slight increase in whole-genome GC content, e.g., from 49.5% to 50.0%, requires thousands of AT to GC mutations, which are unlikely to happen simultaneously. Every single mutation could only change the GC content of one nucleotide among the millions of nucleotides in a prokaryotic genome and result in a tiny change in the double helix stability that is unlikely to be perceived by natural selection.

Upon comparing the correlations between various genomic components, it was observed that the GC content of the third codon position and intergenic sequences exhibited a stronger correlation with halophilicity than that of the first and second codon positions (Table 2). This result indicates a strong correlation in sequences with weaker functional constraints. It was also noticed that the correlations in tRNA and 5S rRNA genes were stronger than those in 16S rRNA and 23S rRNA genes. The tRNA genes are known for their evolutionary flexibility, as they mutate quickly and are sites of genomic instability (49-51). On the other hand, the ribosomal RNA genes are known for their conservation in evolution. However, 5S rRNA genes exhibit higher intragenomic diversity than 16S or 23S (52). In addition, the average substitution rate per million years for 5S rRNA genes is about half of 16S rRNA genes, 0.5% vs. 1% (53). Therefore, the correlations between halophilicity and GC content among structural RNA genes are also significant in sequences with weaker functional constraints. Researches indicate that high concentrations of salt and organic solutes may damage DNA sequences directly or via oxidative stress, inducing DNA double-strand breaks (DSB) (54-56). Consequently, bacteria living in high salt concentrations should have evolved more efficient repairing systems than nonhalophiles. From bacteria to humans, gene conversion acts as an essential role in repairing DSBs (57). Increasing GC content has been documented as part of the recombination process in gene conversion (58-61). Hence, the correlation between GC content and halophilicity is attributed to this nonadaptive process.

No statistically supported conclusion can be drawn for high GC content in halophilic archaea, owing to the limited number of phylogenetic branches diverging in halophilicity. However, almost all the solitary cases, like the class Halobacteria, indicated halophilic archaea have high GC content (11, 18-22). Most halophilic archaea are polyploid and likely use efficient gene conversion to repair their mutations and DSBs (62, 63).

## Conclusions

The commonly observed high GC content in halophilic archaea is widely believed to be an adaptation to avoid UV-induced pyrimidine dimer formation due to their typically high-intensity UV radiation habitats. By pooling data from three comprehensive databases, a dataset of halophilicity was established, including 1226 bacterial species and 181 archaeal species. Phylogenetic correlation analysis using PGLS demonstrated that genome-wide GC content and halophilicity are significantly correlated in bacteria but not archaea. Further analysis of the correlation between halophilicity and GC content among seven genomic components does not support the widespread photoprotection hypothesis or another hypothesis that the high GC content in halophiles stabilizes nucleic acid structures but instead aligns with a nonadaptive hypothesis. Halophiles evolved high GC content probably due to frequent GC-biased gene conversion, which response to the DNA DSBs induced directly or indirectly by the high salt concentrations.

## Methods

The halophilicity data for bacteria and archaea were obtained from three databases: Halodom (32), Protrait (33), and a prokaryotic trait dataset constructed by Madin et al. (34). Among these, the Halodom database, which was manually curated from published literature, was considered the most reliable. The dataset compiled by Madin et al. through integrating multiple databases was considered the next most reliable. The ProTraits database, which employs semi-supervised text mining methods to extract data from online databases and literature, was considered the least reliable. We consolidated the data from these three sources, prioritizing the Halodom database in case of discrepancies, followed by the dataset of Madin et al. and then the ProTraits database. In total, we obtained the halophilicity of 2871 prokaryotic species.

We retrieved the phylogenetic tree provided by the TimeTree database (64) and obtained the phylogenetic information of 1226 bacterial species (816 non-halophiles and 410 halophiles) and 181 archaeal species (8 non-halophiles and 173 halophiles).

The GC contents of prokaryotic whole genomes were gathered from the NCBI Genome Database (https://ftp.ncbi.nlm.nih.gov/genomes/GENOME_REPORTS/prokaryotes.txt, accessed on 2022-06-25). In cases with multiple sequenced strains, the median GC content of all strains was utilized to represent the GC_w_ of the respective species.

We downloaded the genomes designated as reference or representative genomes by the NCBI database for each species and re-annotated them using the DFAST software (version 1.2.11) with default parameters (65). We calculated the GC content of different components in the genome, including protein-coding sequences, intergenic regions, tRNA genes, 5S rRNA genes, 16S rRNA genes, and 23S rRNA genes, based on the annotation results from DFAST. For species without a designated reference or representative genome, we annotated all sequenced genomes of that species. We used the median GC content of each component of a species to represent the species data.

The *phytools* package (Version 0.7–70) in R (Version 4.0.3) (66) was used to determine the phylogenetic signals (λ) and to conduct the phylogenetic paired t-tests. The PGLS regressions were conducted using the *phylolm* package (version 2.6.2) in R (Version 4.0.3) with default parameters (67).

In this study, we did not present the Benjamini-Hochberg (BH)-adjusted *P* values for multiple correlation analyses of the same dataset. In most cases, the *P* and adjusted *P* values were generally on the same side of 0.05, smaller or bigger than 0.05. In a few cases where the significance test results changed because of BH corrections (such as the GC_3_ ∼ Halophilicity in Table 2), we focused on the relative strengths of the correlations rather than the absolute correlation significance values.

## List of abbreviations

AIC: Akaike information criterion
BH: Benjamini-Hochberg
BM: Brownian motion model
DSB: double-strand breaks
EB: Early burst model
GC_1+2_: GC content of the first and second codon positions of protein-coding sequences
GC_16S_: GC content of 16S rRNA genes
GC_23S_: GC content of 23S rRNA genes
GC_3_: GC content of the third codon position of protein-coding sequences
GC_5S_: GC content of 5S rRNA genes
GC_inter_: GC content of intergenic sequences
GC_tRNA_: GC content of tRNA genes
GC_w_: Whole-genome GC content
OU: Ornstein-Uhlenbeck model
PGLS: Phylogenetic generalized least squares
PR: Photoreactivity
PR_w_: Whole-genome photoreactivity
PR_p_: Photoreactivity of protein-coding genes
PR_inter_: Photoreactivity of intergenic sequences
PR_tRNA_: Photoreactivity of tRNA genes
PR_5S_: Photoreactivity of 5S rRNA genes
PR_16S_: Photoreactivity of 16S rRNA genes
PR_23S_: Photoreactivity of 23S rRNA genes
UV: Ultraviolet

## Declarations

### Ethics approval and consent to participate

Not applicable.

### Consent for publication

Not applicable.

### Availability of data and materials

The datasets supporting the conclusions of this article are included within the article and its additional file.

### Competing interests

The authors declare that they have no competing interests.

### Funding

This work was supported by the National Natural Science Foundation of China (grant number 31671321). The funders had no role in the design of the study or collection, analysis, and interpretation of data or in writing the manuscript.

### Authors’ contributions

DKN conceived the study. EZH and SS performed the data analysis. DKN and EZH wrote the manuscript. All authors read, improved, and approved the final manuscript.

## Acknowledgements

Not applicable.

## Notes

### Competing Interest Statement

The authors have declared no competing interest.

### Summary of Updates

Some language bugs have been fixed.

